# Parallelized analysis of spatial gene expression patterns by database integration

**DOI:** 10.1101/386086

**Authors:** Daisuke Miyamoto, Hidetoshi Ikeno, Yuko Okamura-Oho, Akira Sato, Teiichi Furuichi, Yoshihiro Okumura, Yoko Yamaguchi, Ryohei Kanzaki

## Abstract

We developed a computational framework for automated integration of a large number of two-dimensional (2D) images with three-dimensional (3D) image datasets located in the standard 3D coordinate. We applied the framework to 2,810 para-sagittal sectioned mouse brain 2D images of *in situ* hybridization (ISH), archived in the BrainTx database (http://www.cdtdb.neuroinf.jp). We registered the ISH images into the mouse standard coordinate space for MR images, Waxholm space (WHS, https://www.nitrc.org/projects/incfwhsmouse) by linearly transforming them into each of a series of para-sagittal MR image slices, and identifying the best-fit slice by calculating the similarity metric value (δ). Transformed 2D images were compared with 3D gene expression image datasets, which were made using a microtomy-based microarray assay system, Transcriptome Tomography, and archived in the ViBrism DB (http://vibrism.neuroinf.jp): the 3D images are located in the WHS.

We first transformed ISH images of 10 regionally expressed genes and compared them to signals of corresponding 3D expression images in ViBrism DB for evaluating the integration schema: two types of data, produced with different modalities and originally located in different dimensions, were successfully compared after enhancing ISH signals against background noise. Then, for the massive transformation of BrainTx database images, we parallelized our framework, using the IPython cluster package, and implemented it on the PC cluster provided for the Brain Atlasing Hackathon activity hosted by Neuroinformatics Japan Center in Japan. We could identify the best-fit positions for all of the ISH images. All programs were made available through the GitHub repository, at the web site of neuroinformatics/bah2016_registration (https://github.com/neuroinformatics/bah2016_registration).

## Introduction

An understanding of anatomical structure and biological function, based on knowledge of gene expression, is crucial for molecular neuroinformatics approaches to complex tissues and organs, such as the mammalian brain. High-resolution magnetic resonance (MR) images of mammalian brains are standardized in three-dimensional (3D) spaces, where anatomical structure images and anatomical areas are visualized and annotated, respectively (Hawrylycz et al. 2011; Johnson et al. 2010; Mazziotta et al. 2001; Papp et al. 2014). MR images are isotropic; thus, using 3D-to-3D computational transformation programs, researchers can transform images of their own experiments into standard-formatted images (Avants et al. 2011).

Gene expression patterns on two-dimensional (2D) surfaces of thin-sliced tissues are detected with microscopic resolution, using a method of *in situ* hybridization (ISH) (Emson 1993). Results of systemic analyses of comprehensive gene expression-anatomy associations, using the ISH method, are archived in the 3D space of the Allen Brain Atlas (Lein et al. 2007). With the exception of this extremely rigorous work, isotropic images are hard to obtain with ISH methods, because of the technical difficulties inherent in preparation of a series of tissue slices—the slices are hard to align along the Z-axis properly and in the sufficiently dense manner required to create images exhibiting the same resolution as the fine X-Y plane images that are obtained by microscopy. Thus, 3D-to-3D transformation programs are not suitable for these images. Consequently, researchers with sparsely aligned 2D sliced-tissue images of their experimental materials must empirically assume the 2D location of the images within the 3D space of their materials. Following this assumption, 2D-to-2D image transformation programs allow better alignment of ISH images to corresponding 2D images, using manual transformation and visual evaluation of alignment (Lee et al. 2010).

In this study, we introduce an automated schema for integration between 2D and 3D image datasets of comprehensive gene expression patterns with the aim of performing ISH image analyses within broader anatomical models of the brain. We have developed parallelized programs for high-speed linear transformation of 2D ISH images to the best aligned position within the 3D standard coordinate space of MR images. To estimate reliability of the transformation results, we have also produced parallelized programs to calculate expression intensities of the transformed ISH images within anatomical areas of the probabilistic atlas located in the standard coordinate space. Then, we compared the calculated intensities to expression intensity data of 3D maps within the coordinate space. In this study, we used three types of image data: (1) Waxholm space (WHS) datasets, which consist of mouse brain MR images along with a probabilistic atlas (Johnson et al. 2010) as standard coordinates for transformation; (2) 2,810 ISH images of the mouse brain, within the BrainTx DB (http://www.cdtdb.neuroinf.jp) (Furuichi et al. 2011; Sato et al. 2008), as 2D ISH images that are transformed into the WHS; and (3) volume data of ViBrism DB gene expression 3D maps that are located within the WHS MR image space (http://vibrism.neuroinf.jp) (Okamura-Oho et al. 2012, 2014). These ISH images exhibit high-resolution information in the X-Y plane, but few images are available for each gene. In contrast, expression maps in the ViBrism DB are generated from volume data of expression density patterns of approximately 37,000 genes/transcripts throughout the 3D brain, though the spatial resolution of these maps is low. Data integration of the two types of gene expression datasets into the standard coordinate system (WHS) is expected to provide better understanding of the spatial gene expression patterns.

## Materials and Methods

### Materials

The datasets analyzed in this study can be found in public repositories, as described in the following three paragraphs.

The Waxholm space (WHS) is the standard coordinate space of the mouse brain (C57BL/6J, male, 8 weeks of age), based on MR images (Johnson et al. 2010). Corresponding datasets include T1, T2, and T2* MR image files, as well as anatomical area mask files of the probabilistic atlas; all files were downloaded from the INCF software center (http://software.incf.org/software/waxholm-space). We computationally produced 256 images by virtual voxel-based sectioning of the WHS mouse brain in the para-sagittal direction, using the mid-line image and a series of left hemisphere images (slices 80–128) as registration targets.

BrainTx DB (http://www.cdtdb.neuroinf.jp) is a database for comprehensive ISH images of genes expressed in developing mouse brains (Furuichi et al. 2011; Sato et al. 2008). We used 2,810 ISH images from the young adult (3 weeks) mouse brain in this study. At this stage, brain structures are matured and gene expression patterns resemble that of the 8 weeks adult brain. Original images were reduced in size to 512 x 256, which was usable for the following registration process.

ViBrism DB (http://vibrism.neuroinf.jp/) datasets are comprehensive gene expression 3D maps of the mouse brain (C57BL/6J, males, from a variety of developmental stages), produced with the microtomy-based method of Transcriptome Tomography: TT (Okamura-Oho et al. 2012). Expression densities in the adult brain (8 weeks) are detected with 36,558 microarray probes for genes/transcripts; the density data are each reconstructed computationally in voxels to create 3D expression maps located within the standard brain coordinate system, WHS. For this study, we selected expression map datasets corresponding to BrainTx ISH images, using NCBI Gene ID (Entrez gene ID) information. The dataset contained ratios of expression volumes in specific anatomical areas to corresponding anatomical area volumes.

## Methods

We developed a registration framework by utilizing an image processing software, Fiji (Schindelin et al., 2012), and image processing library, ITK (Johnson et al., 2015). In the first step, the size of each ISH slice image was adjusted based on the image of the standard brain. We generated a binarized ISH image, which was a slice image where brain regions were filled with the maximum brightness value, 255, using a thresholding operation. The binarized ISH image was registered by affine transformation on a series of midline and hemisphere slice images from the standard brain. The best-matched slice image of the standard brain (matched to the source ISH image) was selected by evaluation of the registration similarity metric value (δ). We obtained the metric value by computing the sum of the Euclidean (L2) norm between transformed ISH and standard brain images (Johnson et al., 2015). The same transformation was applied to original color ISH images to obtain registered ISH images.

The intensity of gene expression, *I(A)*, in each brain region was calculated by the following equation, based on RGB levels in each image pixel.

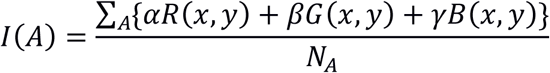

Here, A is a brain region. *R(x, y), G(x, y)*, and *B(x, y)* are pixel values for red, R, green, G, and blue, B, channels located at position x, y within the image. Pixel values were multiplied by the corresponding weight, *α*, *β*, and *γ*, for each pixel in the area A; all multiplication results were added together, then divided by the number of pixels in the region, *N_A_*, to produce *I(A)*. Weights for each color channel, *α*, *β*, and *γ*, were assigned values of −1, −1, and 2, respectively, in our analysis. We enhanced the blue color channel because the most of ISH images contained a large quantity of blue color.

Gene expression densities in 3D maps of the ViBrism DB dataset underwent recalculation to generate expression levels of genes in the corresponding anatomical area of the WHS probabilistic atlas. The levels represented ratios of expression volumes in specific anatomical areas to corresponding anatomical area volumes.

Visual detection of spatial gene expression in the original ISH images was performed by an experimental neurogenetics researcher; highly expressed areas of genes were manually selected from the 39 anatomical areas of the WHS probabilistic atlas.

To calculate these processes on our sets of ISH images, we used the “plato” PC cluster provided by INCF Japan node / NIJC (Neuroinformatics Japan Center), RIKEN; this cluster has four Intel Xeon E5430 CPUs, 16 Intel Xeon X5550 CPUs, and 12 Intel Xeon X5650 CPUs connected to each other by a 40 Gbps InfiniBand network. To execute registration within the cluster, we parallelized our framework using the IPython cluster package. In this scheme, the master node sent an image ID to each slave node, and each slave node independently executed both pre-image and registration processes on the image with this ID. All programs are available through the GitHub repository (https://github.com/neuroinformatics/bah2016_registration).

## Results

### Estimation of the position of the ISH images in the 3D standard space

We designed computational processes for estimation of 3D positions of gene expression, as detected in 2D ISH images (Figure 1). We used WHS images as the standard coordinate, where MR images are provided along with probabilistic atlases of 39 anatomical areas. The estimation workflow was composed of two computational processes: estimation of the 2D ISH image position in the 3D WHS space, and estimation of gene expression intensities in the corresponding anatomical areas. For the first process, we re-sliced the left hemisphere MR image into 49 slices (slices 80–128), on para-sagittal planes, to produce a series of reference sections. Then, we registered the ISH images into each of the sections, using affine transformation. We used a gradient descent optimizer and measured the suitability of each optimization with a similarity metric value (δ) to define the best-fit position of the ISH image within the reference section, on the standard coordinate system (Figure 2). Through use of this method, we found the best-fit slice position on 2,618 of 2,810 images (93%, Supplemental Dataset 1).

**Figure 1.**
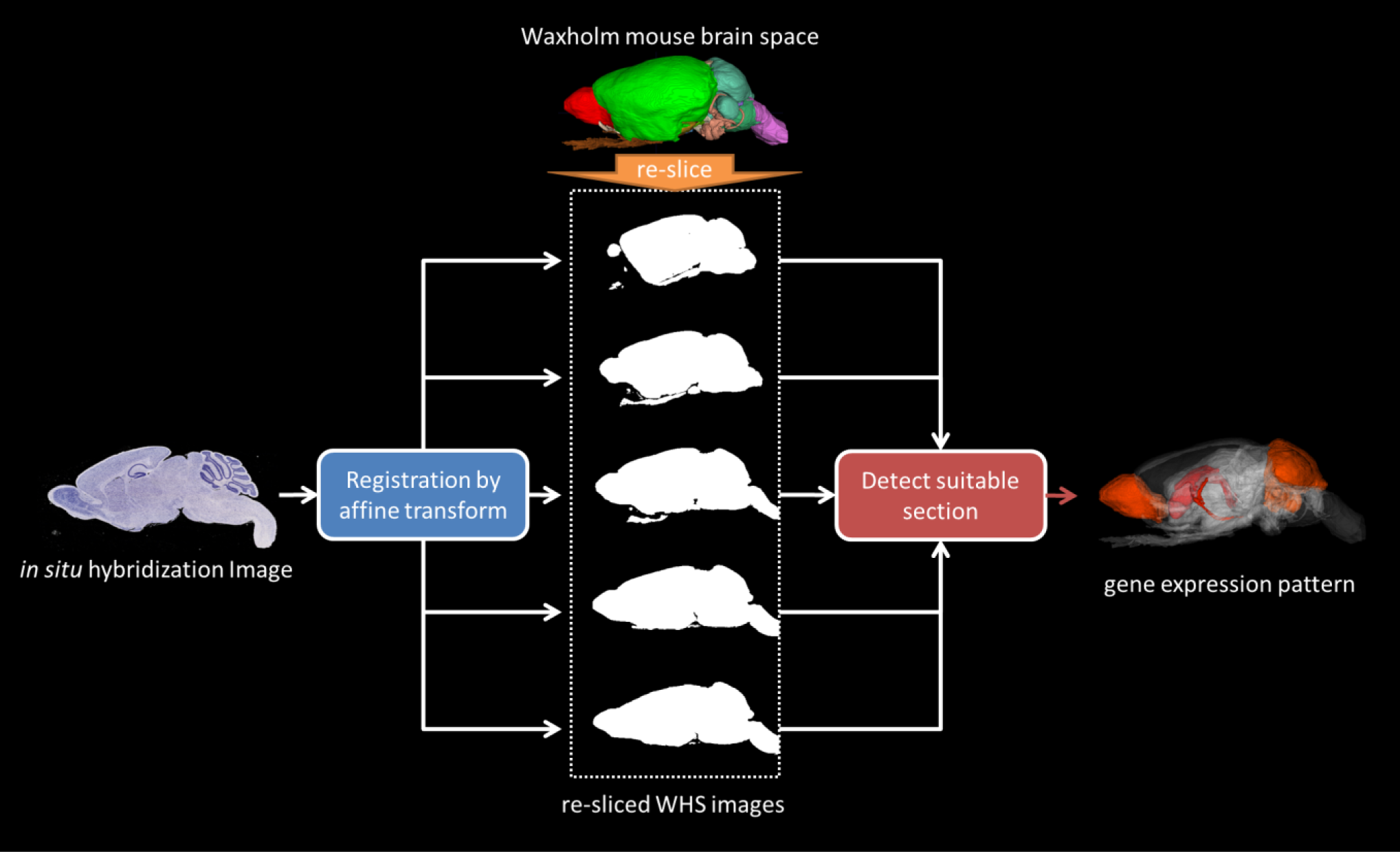
Schema for estimation of three-dimensional (3D) positions of gene expression, as detected in two-dimensional (2D) *in situ* hybridization (ISH) images. The estimation workflow is composed of two computational processes: estimation of positions of 2D *in situ* hybridization (ISH) images within the 3D standard coordinate system of Waxholm mouse brain space (WHS, shown with white arrows) and estimation of regional gene expression intensities in the ISH images, as described using 3D coordinates (shown with a red arrow). The original probabilistic anatomical area atlas is shown with colors in the WHS.

**Figure 2.**
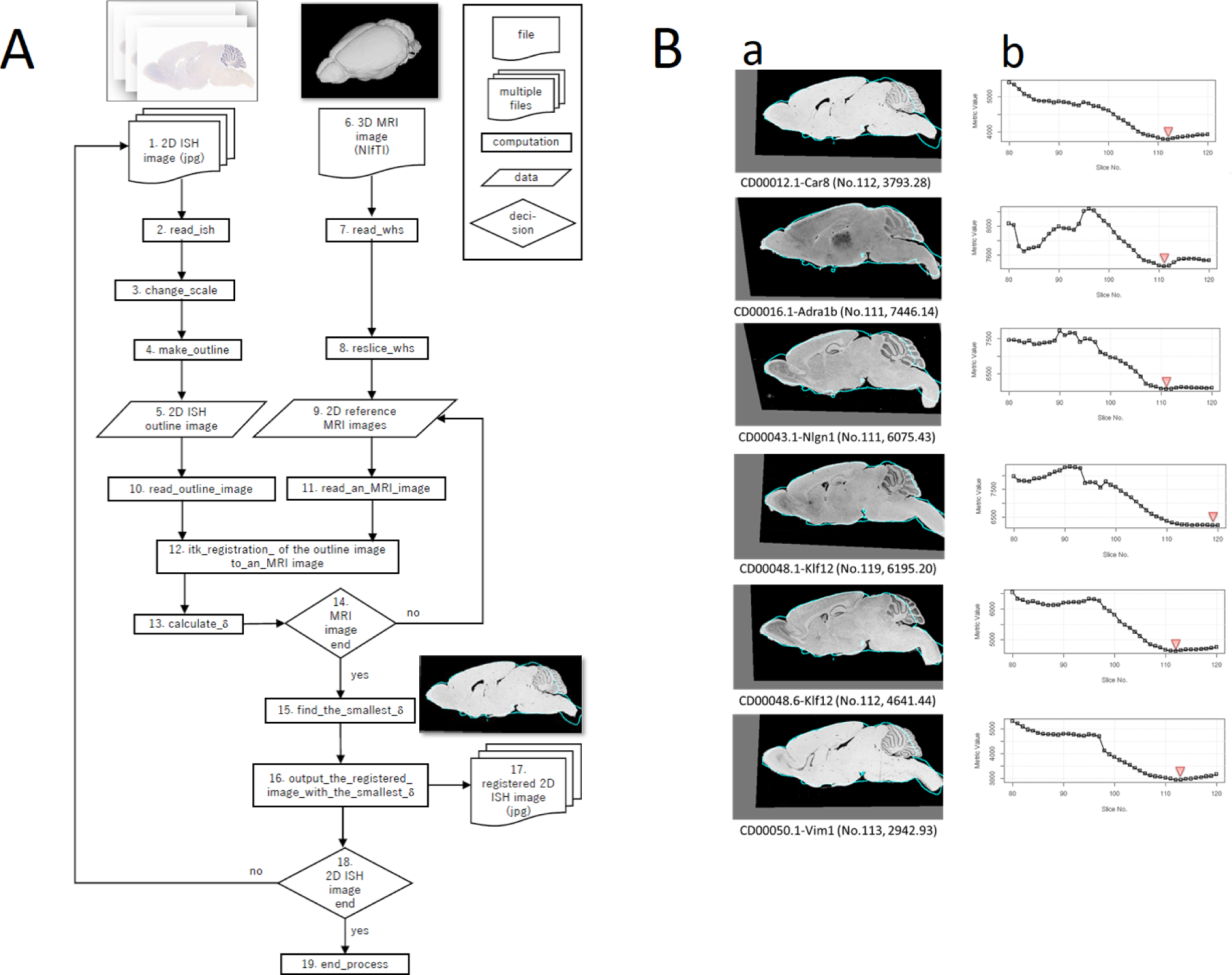
Estimation of positions of two-dimensional (2D) *in situ* hybridization (ISH) images with three-dimensional (3D) standard coordinate system. A: Workflow of the estimation. Processes 1–5: pre-image preparation of 2D images for registration; 6–9: preparation of 3D images for registration; 10–12: registration using affine transformation; 13–19: estimation of the best-fit position of 2D images in the 3D coordinate system. **B. Examples of the best-fit position estimation of six images. a.** 2D ISH images stained with each gene probe and identified as an image ID, as seen below the panel (CD000##. #-gene symbol), are outlined with light-blue lines and transformed to the re-sliced 2D MR para-sagittal images. The best-fit image, estimated as shown in the panel b, is shown with the para-sagittal position number and δ values below the panel. **b.** Metric values (δ) shown in the y axis are calculated in each transformation to the reference MR images, shown in the slice numbers of the x axis. The best-fit position with the smallest δ is indicated with a triangle.

### Estimation of regional gene expression intensity patterns in the WHS anatomical areas

For the second process, color signals of registered ISH images were enhanced with the equation ((Blue × 2.0) – Red - Green) to discriminate ISH-stained signals from background signals (Figure 3A). Reference sections were produced, which included 2D segmentation information of anatomical areas of the WHS brain probabilistic atlas, using the same re-slicing computation as the first process. We evaluated gene expression levels in a variety of anatomical areas by measuring enhanced color signals in the segmented areas of the best-fit reference sections, then calculating regional expression intensities of genes based on average pixel values in each 2D anatomical segmented area. We defined 3D anatomical areas as demonstrating high expression of a given gene if they exhibited high intensity values for the 2D segmented area. Then, we visualized the areas in a map of 3D coordinates. For a test of the estimation processes, we chose genes that are known to be expressed in selected anatomical areas of the mouse brain; we visualized these estimated highly expressed anatomical areas alongside the corresponding MR images (Figure 3B). Our computation schema seemed to successfully estimate regional expression of these genes, based on our initial assessments.

**Figure 3.**
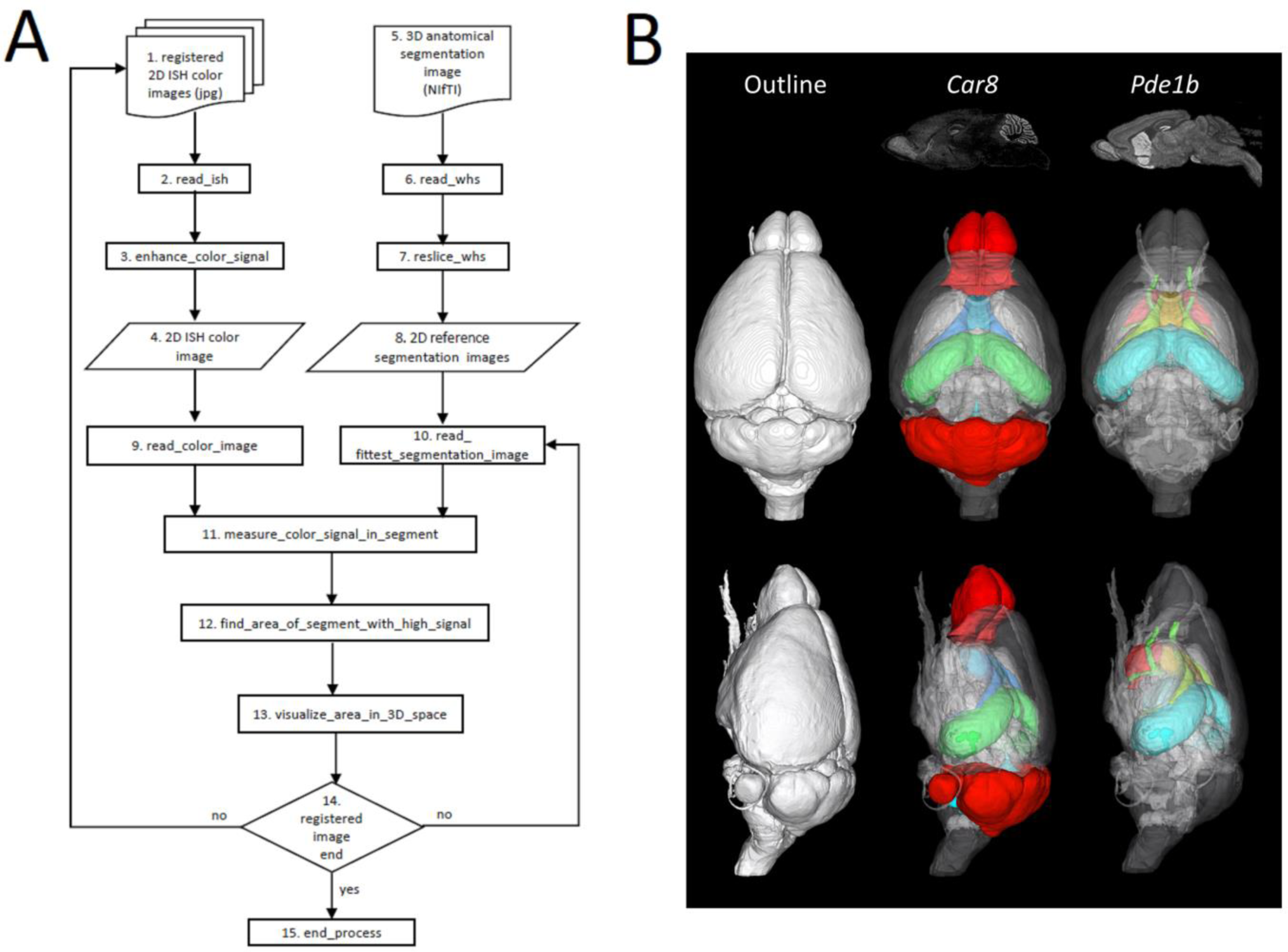
Estimation of spatial gene expression of *in situ* hybridization (ISH) images in the three-dimensional (3D) coordinate system. A: Workflow of the estimation. Processes 1–4: enhancement of color signals; 5–8: preparation of 3D images for registration; 9–11: measurement of color signals of ISH expression densities in segmented anatomical areas; 12–15: visualization of highly expressed anatomical areas in the 3D coordinate space. **B: Visualization of spatial gene expression.** Genes known to be regionally expressed, Car8 and Pde1b, were registered to WHS MR images and color signals were enhanced, as indicated in the first row of the panel. Expression densities were measured in corresponding anatomical areas of the WHS probabilistic atlas and visualized with colors, as seen in the second row. Car8 shows the highest expression in the olfactory bulbs and cerebellum (indicated in red), moderate expression in the hippocampus (in light green), and lowest expression in a part of the cerebral cortex and septal areas (in light blue). Pde1b shows high expression in the nucleus accumbens (in red), striatum (in orange), internal septal nuclei (in yellow), hippocampus (in light blue) and subcommissural organ (light green).

### Comparison of the regional expression intensities to ViBrism DB datasets

We compared spatial gene expression among data from our automated detection method using transformed 2D ISH images, data from visual detection using the original ISH images, and data from recalculation of volume data from the ViBrism DB in the WHS probabilistic anatomical atlas. As is shown in 10 representative regionally expressed genes, >80% of the anatomical areas that exhibited visually recognizable high expression were detected by our automated method as high-expression anatomical areas (Figure 4). Moreover, we found five areas of high expression that were present in computational results of our automated method and in the ViBrism DB recalculated data, but were not present within visual recognition data. Although the computational methods tended to overestimate expression areas (as will be discussed in the Discussion section), the 2D-to-3D registration could enable proper anatomical annotation of areas with complex structures.

**Figure 4.**
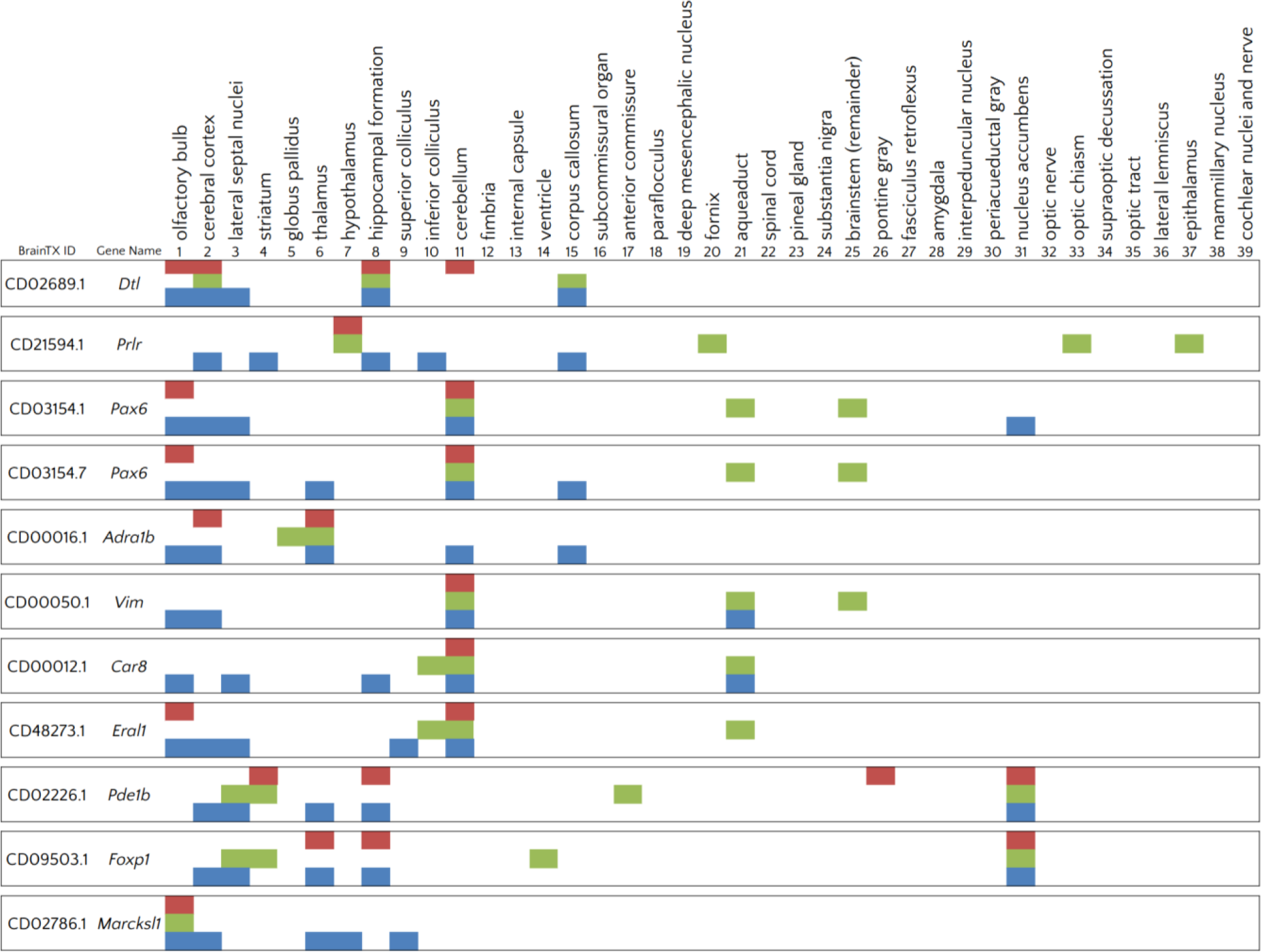
Comparison of estimated spatial gene expression areas. Shown are results of manual detection of histological stains using the original *in situ* hybridization (ISH) images (red boxes), results of computational recalculation of volume data of ViBrism DB (green boxes), and results of automated detection using our framework for two-dimensional (2D) ISH image transformation (blue boxes). Ten selected genes are shown (pax6 was detected with two probes). For nine genes, 10 total areas are commonly recognized as highly expressed areas in all three sets of results. For five genes, five total areas are recognized as highly expressed areas in both computational methods, but not in the manual detection method. Highly expressed areas detected by a single method may be related to false estimation, as discussed in the Discussion section.

### Parallelization of the estimation and comparison processes

We parallelized the computation processes of the estimation of the best-fit position and the subsequent comparisons, with the aim of applying these processes to a wide range of ISH images. The registration process requires an exceptional length of computation time. We estimated the time for applying our method to all 2,810 ISH images in the BrainTx DB; it would require >2 months when using a single computer (Intel Xeon E5410). Therefore, we parallelized the method, using the map method from the IPython parallel package. IPython engines in each calculation node were executed by the Sun Grid Engine (SGE) and received different IDs for ISH images. We completed 2,810 registrations in 1.2 days, using the “plato” PC cluster operated by INCF J-node / RIKEN NIJC (Neuroinformatics Japan Center). This result was 60-fold faster than the estimated single processor execution (Figure 5). All estimated gene expression intensities of ISH images were supplemented (Supplemental Dataset 2).

**Figure 5.**
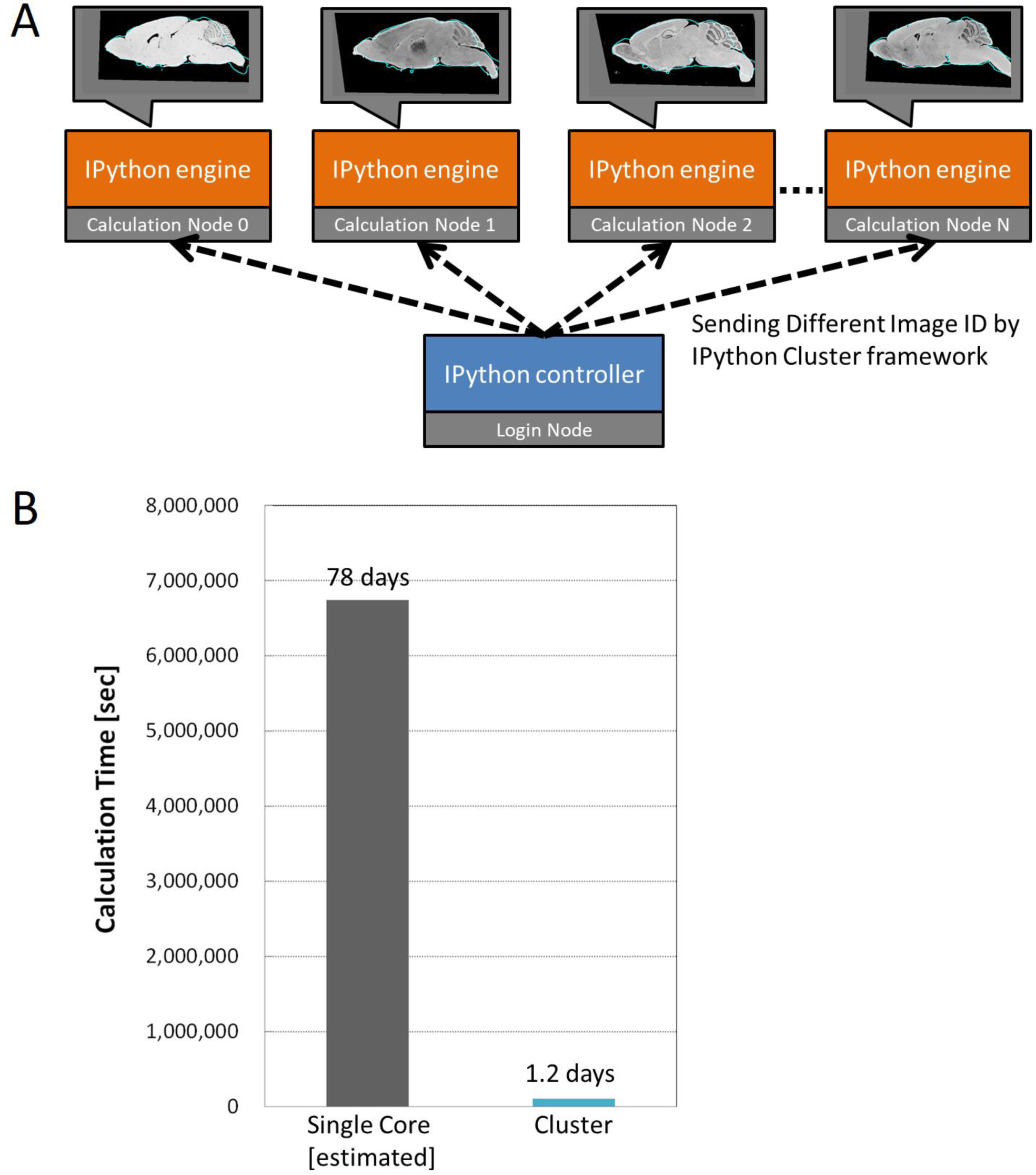
Parallelization methods and results. A: Schematics of parallelization method. We used IPython Cluster for parallelizing registration tasks. Each calculation node has a IPython engine that is invoked by MPI. Each node receives the ID of *in situ* hybridization (ISH) images and calls ITK for registration function. **B**: Calculation time of single CPU processing (estimated) and parallelized processing for 2810 ISH images. Our parallelized scheme achieved 60-fold faster processing than a single CPU, using 152 CPU cores in our cluster (plato).

## Discussion

### Automated registration framework

We developed an automated and massive registration framework for transforming ISH slice images into the 3D standard brain workspace. Identifying the location of an experimentally obtained 2D tissue slice image within the 3D anatomical standard space is generally not very easy; consequently, the identification process requires time and results may differ among multiple researchers. Thus, computational aids are valuable as open resources to facilitate time-conscious image processing and to counterbalance individual researcher differences in identification perspectives.

Various image processing software and libraries have been already developed for registration. We aimed to provide the computational aid for image identification through a combination of these software tools. Our framework was composed of a combination of Fiji macros for pre-image processing, including contrast change, image size change, and a registration program using ITK. A small Python script connected each process and controlled the process pipeline.

### Parallel processing of registration for massive image data

Our framework was parallelized within a PC cluster; this acceleration provided a new scheme for large-scale analysis of gene expression. However, the acceleration ratio (60-fold faster) did not correspond to the number of CPU cores (152 cores in 32 CPUs) within the cluster. We considered that the loading time for each ISH image from NFS-mounted storage was the primary bottleneck of this process. Using multi-layer parallelization—where each CPU processes one image at a time and each core in a CPU processes each slice—could reduce the peak communication time and improve parallelization performance. Alternative methods to improve performance include use of a GPU; notably, there have been some implementations of a GPU registration method (Shamonin et al., 2013). Joint use of large-scale parallelization and GPU registration could provide a method for easy handling of a large amount of image data.

### Registration of ISH image and its evaluation

In this study, 93% of ISH images were selected as the appropriate slice and registered properly into the 3D brain map. However, 192 ISH images could not be registered by our framework. These failed cases of registration were detected by transformation of the resulting image and changes in the similarity metric values. In most failed cases, the similarity metric value was oscillatory, and changed at several local minima by increasing the slice number.

Failed registration could be caused by several factors, including lack of information for registration or large individual differences in brain shape. In our registration framework, only the outline of brain shape was used to evaluate the fitness of brain shape. We assumed that all slice images were made by cutting the brain perpendicularly, but this approach might prove to be very difficult in a research setting. Thus, our framework could be improved by considering the tilt of the slicing operation.

Furthermore, the registration process could be improved by considering the position and shape of brain regions. We can apply more flexible and precise translation, such as thin-plate spline, when fitting ISH images to reference slice images, given sufficient image characteristics.

### Estimation of gene expression area and its evaluation

We estimated intensities of gene expression from registered ISH images and compared them to volume images of the ViBrism DB that were produced with the TT method. Most visually recognizable gene expressions were also recognizable within our automated framework as high-expression anatomical areas. Moreover, some areas of undetectable gene expression were found with our framework and confirmed with ViBrism DB data. Example analyses are shown in Figure 4.

Exact estimation of gene expression intensity is difficult because pixel colors of ISH images are affected by various factors, including background staining, cell densities and sizes within ISH tissues, mRNA structures and densities, and probe qualities within hybridization experiments. We computationally attempted to minimize background staining interference and enhance signals with the blue-color enhancing method. Estimation accuracy may be affected within image registration methods, as discussed in the previous section. Registration programs using non-linear transformation, as well as transformation to arbitrary planes, need to be improved for better estimation. Advanced registration programs would be helpful in reducing inappropriate estimation of expression areas, particularly for genes expressed in small anatomical areas. Our automated framework seems effective for estimating outlines of gene expression areas and visualizing them alongside detailed images from the WHS anatomical atlas.

Integration of information obtained from images in a variety of modalities could be important for better understanding of gene expression patterns and intensities, along with the underlying biology. Here, we integrated 2D ISH images of gene expression into the 3D MRI space and compared these integrated images with other volume data that describe gene expression, within ViBrism DB datasets. Indeed, overestimation occurs in gene expression areas contained within the ViBrism DB datasets, as a result of the TT method (described in a previous paper) (Okamura-Oho et al. 2012). Comparison of the two results could help to identify falsely positive expression areas within the 3D maps.

Nevertheless, overlying transformed 2D ISH images, which precisely show local cell-level gene expression, and ViBrism DB 3D maps, which provide an overview of the relative intensities of comprehensive gene expression throughout the brain on WHS MR images, may provide researchers with exclusive information regarding functional anatomy associations within the brain. Thus, we are developing a web-based service platform to browse the data discussed in this report.

### Hackathon activities for collaboration of experimentalists and computer scientists

Hackathon activities were performed effectively for interdisciplinary collaboration to create new data analysis tools. Our registration framework was efficiently invented during the hackathon organized by INCF J-node/ RIKEN NIJC (https://www.neuroinf.jp/bah2015/). In 2015, we developed a prototype of our framework in the first NIJC hackathon (Brain Atlasing Hackathon), by collaboration with experimentalists and programmers. In 2016, we developed the parallelized version, for running on the cluster machine, during the second hackathon. We expect that our newly developed programs will serve as a valuable open resource to improve the utility of ISH images in studies of whole brain gene expression.

### Information Sharing Statement

All programs for integration of 2D/3D images were made available through the GitHub repository, at the web site of neuroinformatics/bah2016_registration (https://github.com/neuroinformatics/bah2016_registration). Original images of 2,810 para-sagittal sectioned mouse brain 2D ISH images are browsable in the BrainTx database (http://www.cdtdb.neuroinf.jp), MR images of WHS are downloadable from the INCF/NITRIC website (https://www.nitrc.org/projects/incfwhsmouse) and 3D gene expression image datasets are archived in the ViBrism DB (https://vibrism.neuroinf.jp/search.html). Integrated results of 2D ISH images and 3D gene expression images in MR images of WHS are browsable and shareable using URLs of the ViBrism DB (example: Adra1b gene https://vibrism.neuroinf.jp/setsearch/3d/view/Cx1/1a32c51a5e001bdc64c7365be238546c).

## Acknowledgements

A part of this work was supported by RIKEN Neuroinformatics Japan Center (NIJC). WHS MR images were provided from INCF Waxholm Space Task Force of the Program on Digital Brain Atlasing. This work was mainly performed in Brain Atlas Ideathon/Hackathon and NIJC Hackathon organized by NIJC.

BrainTx and ViBrism DB are supported by the grant from RIKEN Neuroinformatics Japan Center. ISH image production of BrainTx was supported by JSPS KAKENHI 23300137 and 258057 to TF. ViBrism DB was supported by JSPS KAKENHI 25560428, 26280110, 268032, 15HP8038, 16HP8032, and 17HP8082; and the RIKEN Strategic Programs for R&D to YO. Development of the program was supported by Grant-in-Aid for JSPS Fellows 16J09788 to DM and JSPS KAKENHI 25330342 to HI.

## Compliance with ethical standards

All procedures involving animals and their care to create datasets of BrainTx and ViBrism DB were carried out in accordance with the recommendations of the RIKEN Regulations for Animal Experiments, as described previously (Furuichi et al., 2011; Sato et al., 2008, Okamura-Oho et al., 2012).

## Conflict of Interest

The authors declare that the research was conducted in the absence of any commercial or financial relationships that could be construed as a potential conflict of interest.

## Author Contributions

DM, HI, and YO designed the workflow. DM and HI developed the program. DM, Yok, and YY parallelized the program. YO, AS, TF, and RK managed and processed databases. All authors wrote the manuscript.

## Supplementary Material

### Supplemental Dataset 1. Metric values of the registration in each ISH image

This contains 2,810 text files. Each file is named with CD ID that is a 2D ISH file identification in the BrainTx database. The text in a CSV format shows slice numbers of para-sagittal sections (80-128) in the first column and similarity metric values (δ) in the second column.

### Supplemental Dataset 2. Estimated gene expression intensity from ISH images

This dataset shows estimated gene expression intensities of the 2,810 ISH images of the BrainTx database in the 39 anatomical areas of the WHS brain probabilistic atlas. The text in a CSV format shows CD IDs of ISH images in the first column, background intensities in the second column and estimated intensities of the areas in the following 39 columns. The column numbers represent anatomical areas shown in Figure 4.

